# Natural Selection of the Common *STING* Allele *HAQ* in Anatomically Modern Humans

**DOI:** 10.64898/2026.09.04.749500

**Authors:** Alexandra Aybar-Torres, Lei Jin

## Abstract

The *STING* allele R71H-G230A-R293Q (*HAQ*) was positively selected in Anatomically Modern Humans (AMH) during the Out of Africa (OoA) migration. Here, we show that 51.61% of Native Americans (NAM) are *HAQ/HAQ*, while only 6.45% are *WT/WT. HAQ* individuals are defective in DNA-induced type I IFNs responses, such as smallpox-induced IFNα production. 82.3% of NAM carry the *HAQ* allele, which helps explain the post-Columbus NAM population collapse from the smallpox epidemic. Similarly, in 12 sub-groups of Sub-Saharan Africans, we found 0% *HAQ/HAQ*, indicating ongoing negative selection reflecting Sub-Saharan Africa as the world’s heaviest infectious-disease regions. To understand the positive selection of *HAQ* in non-Africans, we dephased *HAQ* haplotypes in 74 sub-populations worldwide, over ~4,000 individuals. We found that the *HAQ* allele evolved faster than the neutral *H232 STING* allele and had more generations during the OoA migration, increasing its frequency in non-Africans. A mouse model of *HAQ* housed in a Specific Pathogen-Free mouse facility generates more surviving pups than the *H232* mice. In summary, the *HAQ* allele was positively selected during OoA migration but is negatively selected in Sub-Saharan Africa, underscoring its essential role in AMH adaptation to diverse environments.

## Introduction

Genetic data from genome-wide (*1, 2*), mitochondrial DNA (*3*), and Y-chromosomal analyses (*4*) establish that Anatomically modern humans (AMH) outside of Africa are descendants of the Great Out-of-Africa Migration (OoA) 50,000~70,000 years ago. However, very little is known about the natural selection events during the OoA migration. For example, though it can be expected that the adaptations during the OoA migration would have driven a beneficial allele to high frequency, modern human populations globally show few classical genetic signatures of strong selection (*5, 6*). Similarly, although negative selection is one of the most important mechanisms of evolution and plays a crucial role in maintaining the long-term stability of biological structures by removing deleterious mutations, no negatively selected alleles have been found during OoA migration. Key questions in OoA migration remain: Was there a natural selection event, either positive or negative selection, during the OoA migration? If so, which people were selected and flourished? And who were the people extinguished? What were the selective factor(s)? What functions of the genes were positively or negatively selected? Is the natural selection event happening now?

STING (stimulator of interferon genes) is an essential player in cytosolic DNA-induced type I IFN responses (*7, 8*). The DNA-cGAS-cyclic dinucleotides (CDNs)-STING-Type I IFNs pathway is a central innate immune sensing mechanism that detects cytosolic double-stranded DNA (from pathogens, damaged mitochondria, or nuclear leakage) as a danger signal. Upon binding DNA, the enzyme cyclic GMP-AMP synthase (cGAS) activates and catalyzes the synthesis of the second messenger 2′3′-cyclic GMP-AMP (cGAMP) from ATP and GTP. This cyclic dinucleotide then binds to and activates the endoplasmic reticulum (ER)-resident protein STING, triggering its oligomerization and translocation to the Golgi. There, STING recruits and activates the kinase TBK1, which phosphorylates the transcription factor IRF3; phosphorylated IRF3 dimerizes, enters the nucleus, and drives robust transcription of Type I interferons. These interferons establish an antiviral state, promote adaptive immunity, and contribute to inflammation and antitumor responses, making the pathway a key regulator of host defense and a therapeutic target in infection, autoimmunity, and cancer. The 2024 Albert Lasker Basic Medical Research Award was awarded for the discovery of the cGAS enzyme that senses foreign/self DNA and stimulates immune and inflammatory responses.

In 2011, we discovered the common human *STING* allele *HAQ*, which contains three nonsynonymous SNPs (R71<u>H</u>, G230<u>A</u>, and R293<u>Q</u>) (*9*). We found that *HAQ* is the 2^nd^ most common *STING* allele in Americans, but is defective in type I IFNs stimulation (*9, 10*). In 2022, we discovered that the *HAQ* allele was positively selected, while its ancestor allele *AQ* was negatively selected during the OoA migration (*11*). ~1.4% of Sub-Saharan Africans have the *HAQ* allele, while ~63.9% of East Asians are *HAQ* carriers (*11*). ~40.1% of Sub-Africans are *AQ* carriers, while ~0.4% of East Asians have the *AQ* allele (*11*). Notably, the *HAQ* and *AQ STING* alleles have a dominant-negative effect because STING exists as a homodimer (*9, 12-14*). Here, we investigate *HAQ* evolution during OoA migration to reveal the cause of *HAQ* selection, its historical impact, and current implications.

Controversy remains over whether the *HAQ* human *STING* allele differs functionally from the *WT* allele (*15, 16*). We generated an *HAQ* mouse model (10, 14, 15), conducted an *HAQ* clinical trial (NCT02471014) (17), and performed *HAQ* human genetic analysis in the 1000 Genome Project (1KGP) and the Alzheimer’s Disease Sequencing Project *(11, 17)* to show that *HAQ* is functionally different from the *WT STING. HAQ* humans have different susceptibility to human diseases, including infectious and inflammatory diseases and cancer (*18-22*). Here, we further show that *HAQ* was positively selected, while *WT* was negatively selected, in NAM (Figure 1). However, in 2017, a group from Aduro Biotech claimed that “*the HAQ allele, far from being null, is functionally responsive and do not expect the outcome or interpretation of clinical trials targeting STING will be impacted by inclusion of patients bearing the HAQ allele*” (*16*). STING clinical trials all failed (NCT03010176, NCT02675439) (*23, 24*). A recent Phase I clinical trial of a STING agonist (XMT-2056) resulted in a fatal adverse event (NCT05514717). Nevertheless, several groups continue to argue that the *HAQ* is functionally similar to the *WT STING* alleles (*25-27*). This is because if *HAQ*, the 2^nd^ most common human *STING*, is defective in DNA sensing and type I IFNs production, as we showed before (*9-11*) and here by the identification of purifying selection of *HAQ* in Sub-Saharan Africans, then the DNA-cGAS-CDNs-STING/HAQ-Type I IFNs is non-functional in *HAQ* individuals, which is 82% of Native Americans (NAM), ~65% of East Asians (EAS), and ~42% of South Asians (SAS). Furthermore, the positive selection of *HAQ* will render the DNA-cGAS-STING pathway dispensable and obsolete in non-Africans. Instead, STING’s DNA-sensing-independent function in human reproduction was positively selected.

**Figure 1.**
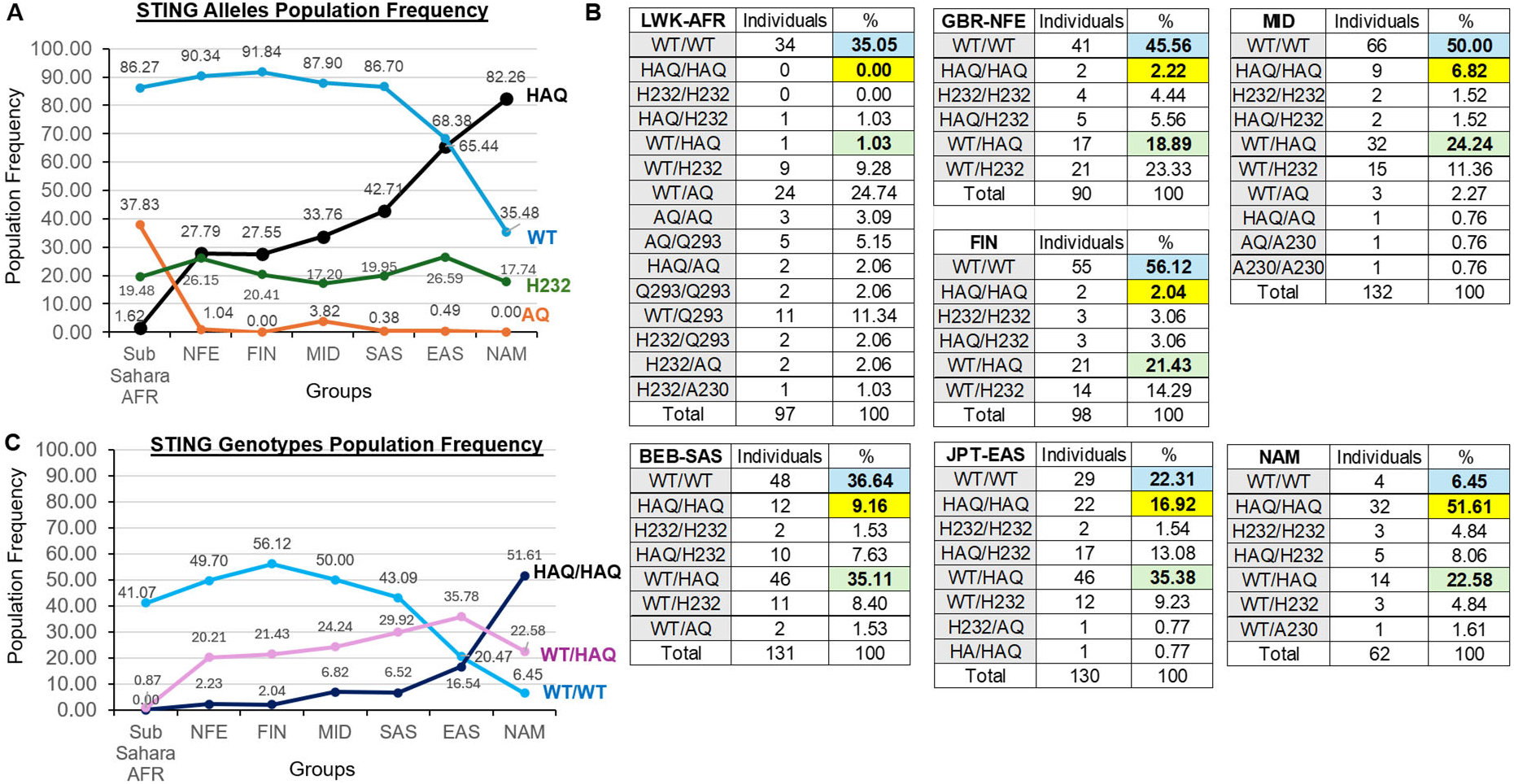
The *HAQ* allele was positively selected during the OoA migration. **A**. Percentage of people who carry the indicated *STING* allele in the populations. Genotype data from Sub-Saharan AFR, Middle East, Non-Finnish Europeans, Finnish, South Asians, East Asians, and Native American populations were from the 1000 Genomes Project (1KGP) and the Human Genome Diversity Project (HGDP). **B**. Human *STING* genotypes and their frequencies in the indicated groups from 1KGP+HGDP. **C**. Population frequencies of *WT/WT, WT/HAQ*, and *HAQ/HAQ* in the seven major populations in 1KGP and HGDP.

## Results

### 51.61% of Native Americans are *HAQ/HAQ*, while just 6.45% are *WT/WT*

Analyzing data from 1KGP, we previously showed that the *HAQ* allele was positively selected during the OoA (*11*). In comparison, the *H232* allele is neutral (*11*). The 1KGP does not include groups representing Native Americans (NAM) or Middle Easterners (MID), two key groups in OoA migration. Here, we analyzed the Human Genome Diversity Project (HGDP) (~828 individuals, 54 groups). The HGDP contains 5 NAM groups: Colombian, Karitiana, Maya, Pima, Surui, and 3 MID groups: Bedouin, Druze, Palestinian. We extracted high-quality individual-level *STING* genotypes from the gnomAD v3.1.2 genomes dataset. We obtained the data using a cloud-based pipeline on Google Cloud Platform (Materials and Methods). We then combined individual-level *STING* genotypes from HGDP and 1KGP (~4,000 individuals, 74 groups) and analyzed them in Figure 1.

In the seven major super-groups: Sub-Saharan African (AFR), Non-Finnish European (NFE), Finnish (FIN), MID, SAS, EAS, and NAM, the population frequency of individuals who carry the *HAQ* allele (including *WT/HAQ, HAQ/HAQ, HAQ/H232*, et al.) showed a clear cline consistent with positive selection during OoA migration (Figure 1A, 1B). In AFR, ~1.62% of people carry the *HAQ* allele, whereas ~82.26% of NAM are *HAQ* (Figure 1A). MID group exhibited intermediate values consistent with their transitional place during OoA migration. Genotype-level analysis further supported this pattern (Figure 1C).

Homozygous *HAQ* (*HAQ/HAQ*) individuals were absent (0 out of 801 individuals across 12 Sub-Saharan African groups, but 51.61% of NAM are *HAQ/HAQ* (Fig 1C). Only 6.45% of NAM are *WT/WT* (Figure 1C). The sharp rise in *HAQ* allele frequencies outside Africa, coupled with near-fixation in NAM and the maintenance of high haplotype diversity, is consistent with positive selection on the *HAQ* allele during OoA migration. Next, we calculated the selection coefficient (s) for positive selection on the *HAQ* allele.

### The *HAQ* allele underwent a moderate, continuous positive selection during the OoA migration

The *HAQ* allele displayed a pronounced frequency cline across global populations (Figure 1). In AFR, the allele occurred at low frequency (1.62%), consistent with its ancestral state. Frequencies increased progressively in non-African groups, reaching 27.8% in NFE, 33.8% in MID, 42.7% in SAS, 65.4% in EAS, and 82.3% in NAM (Figure 1A). This corresponds to a 17-to 51-fold elevation relative to the African baseline, with the most dramatic rises observed along the East Eurasian and NAM branches (Table 1). On the logit scale, this reflects a substantial increase in allelic odds: approximately 115-fold higher in EAS and 281-fold higher in NAM compared to the African proxy (Table 1). By comparison, the neutral control *H232* allele exhibited remarkable stability (mean 21.1 ± 3.5% SD across non-African populations; range 17.2–26.6%), indicating that the cline in the *HAQ* allele is not attributable to demographic history or sampling artifacts alone.

**Table 1.**
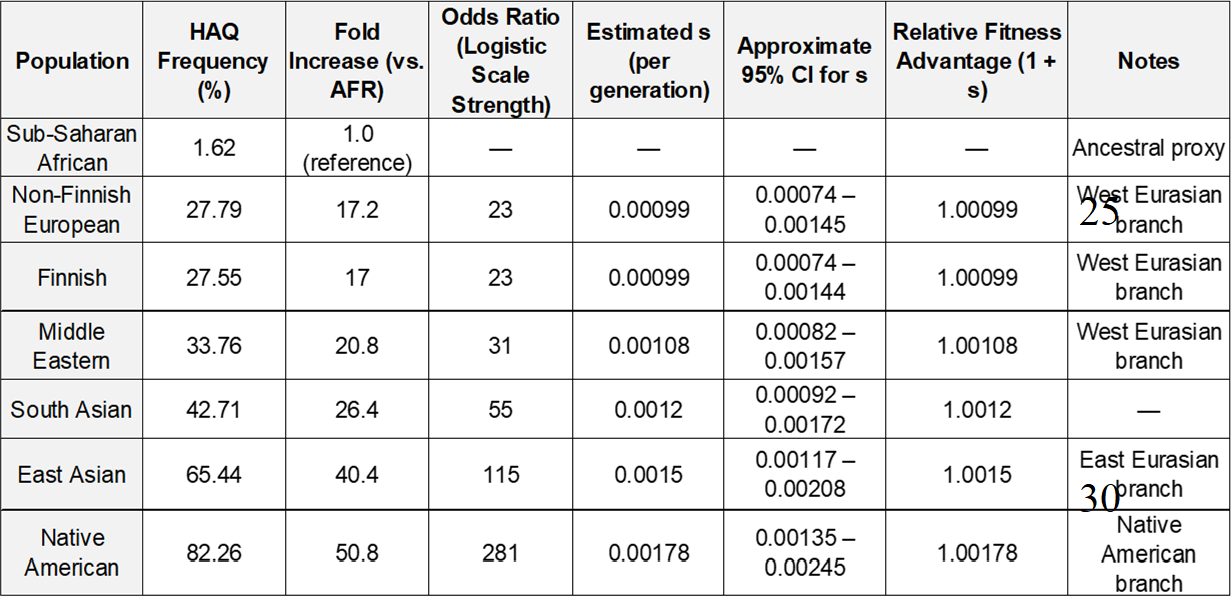
The HAQ allele was under moderate, continuous positive selection during the OoA migration. Selection coefficient (s) was estimated using the deterministic logistic model: s ≈ [ln(p_1_/(1−p_1_)) − ln(p_0_/(1−p_0_))] / t, where p_0_ = African frequency, p_1_ = target population frequency, and t = generations since Out-of-Africa (~3,171 using G = 20.5 years). 95% CIs derived from Monte Carlo simulation incorporating uncertainty in p_0_, modern frequencies, and generation time/divergence dates. Branch-specific estimates highlight stronger effective selection on the East Eurasian → Native American lineage.

Under a deterministic logistic selection model calibrated to ~65,000 years since the OoA dispersal (~3,171 generations at an ancestral generation time of 20.5 years in non-Africans (*28, 29*)), we estimated a moderate selection coefficient of s = 0.00150 per generation for the lineage leading to East Asians (95% CI 0.00117–0.00208, derived from Monte Carlo sensitivity analysis incorporating uncertainty in divergence times, generation intervals, and starting frequency) (Table 1). Branch-specific estimates revealed weaker effective selection on the West Eurasian branch (s ≈ 0.00099; 95% CI 0.00074–0.00145) but stronger selection on the East Eurasian-to-NAM lineage (s ≈ 0.00178 for NAM overall; up to ~0.0020–0.0028 on the terminal ~22 kya branch) (Table 1). Collectively, these results demonstrate that sustained, moderate positive selection acting over deep time drove the substantial rise in *HAQ* frequency in non-African populations.

### *HAQ/HAQ* is negatively selected in AFR

To understand the evolution of *HAQ* haplotypes, we previously dephased all *HAQ* and *AQ* genotypes in the LWK (East Africans) and CHS (Southern Chinese) groups based on 19 SNPs (MAF>1%) in the 18.5kb region covering the human *STING* gene (*11*). The human *STING* gene is ~7.2kb long. We detected 10 *AQ*, 2 *HAQ* haplotypes in LWK, and 6 *HAQ*, 0 *AQ* sub-haplotypes in the CHS population (*11*). We further found that i) *HAQ-1* is dominant with a 34.76% haplotype population frequency in CHS; ii) *HAQ-2* was derived from the *AQ-1* and is the founder *HAQ* haplotype (*11*). The low genetic diversity of the *HAQ* haplotype in LWK, compared with CHS, contradicts the notion that AMH genetic diversity is substantially higher in African populations than in populations outside Africa, one of the most consistent and well-supported findings in human population genetics. African genomes typically contain nearly a million more variants than non-African genomes on average. The lack of genetic diversity of the *HAQ* haplotype in the LWK group suggests that *HAQ* may be negatively selected in sub-Saharan Africa. To test this hypothesis, we extracted all the *HAQ* and *AQ* haplotypes from ~4,000 individuals in 74 groups in 1KGP+HGDP.

We identified 22 *HAQ* haplotypes and 22 *AQ* haplotypes (Table S1, S2). The non-Africans have all 22 *HAQ* haplotypes (Figure 2A). However, AFR has only *HAQ-1* and *HAQ-2*. This confirms that AFR populations lack *HAQ* genetic diversity. The PopArt map showed that *HAQ-2* is 1 nucleotide away from *AQ-1* (Figure 2B and 2C). Notably, *HAQ-2* exists in the San and Pygmies hunter-gatherer AFR groups (Figure 2D). The San and Pygmies groups separated from other AFR groups 100 ~200kya ago, before the OoA migration (*30-32*). Thus, *HAQ-2* has been in AFR for at least 100ky, but there was no genetic drift from *HAQ-2*. In addition, there is no homozygous *HAQ* individual (*HAQ/HAQ*) in any of the 12 AFR groups in 1KGP+HGDP (Figure 2E). In comparison, in 2 indigenous groups in the Amazon rainforest, 18 out of 20 are *HAQ/HAQ* (Figure 2E). In the Papuan highlanders, 12 out of 17 are *HAQ/HAQ* (Figure 2E). The Papuan highlanders have been isolated for at least 50,000 years (*33-37*). The Amazon groups have been isolated for 10,000 years (*38*). The S (selection coefficient) of *HAQ/HAQ* in Sub-Saharan Africa is −0.00138 (Table 2), indicating a moderate purifying selection. Amazonian groups show strong recent positive selection for *HAQ/HAQ* (s = 0.00437) (Table 2).

**Table 2.**
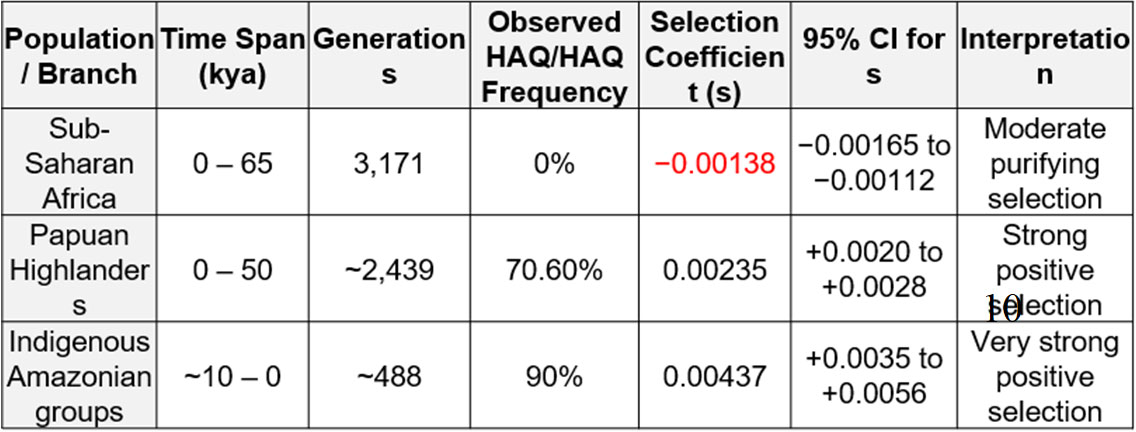
*HAQ/HAQ* is negatively selected in Sub-Saharan Africans. The table summarizes the branch-specific selection coefficients estimated using deterministic logistic forward simulations over 3,171 generations (~65,000 years at 20.5 years per generation). 95% confidence intervals for s were derived from Monte Carlo simulations (varying ancestral starting frequency between 0.5–1.2%) combined with 2,000 stochastic Wright–Fisher replicates per condition (N_e_ = 10,000–50,000). Under purifying selection in Africa (s = −0.00138), *HAQ/HAQ* homozygotes are consistently driven to near-zero frequency. In contrast, positive selection outside Africa was substantially stronger, with branch-specific coefficients ranging from s = +0.00235 on Papuan Highlanders to s = +0.00437 on the Amazonian terminal branch.

**Figure 2.**
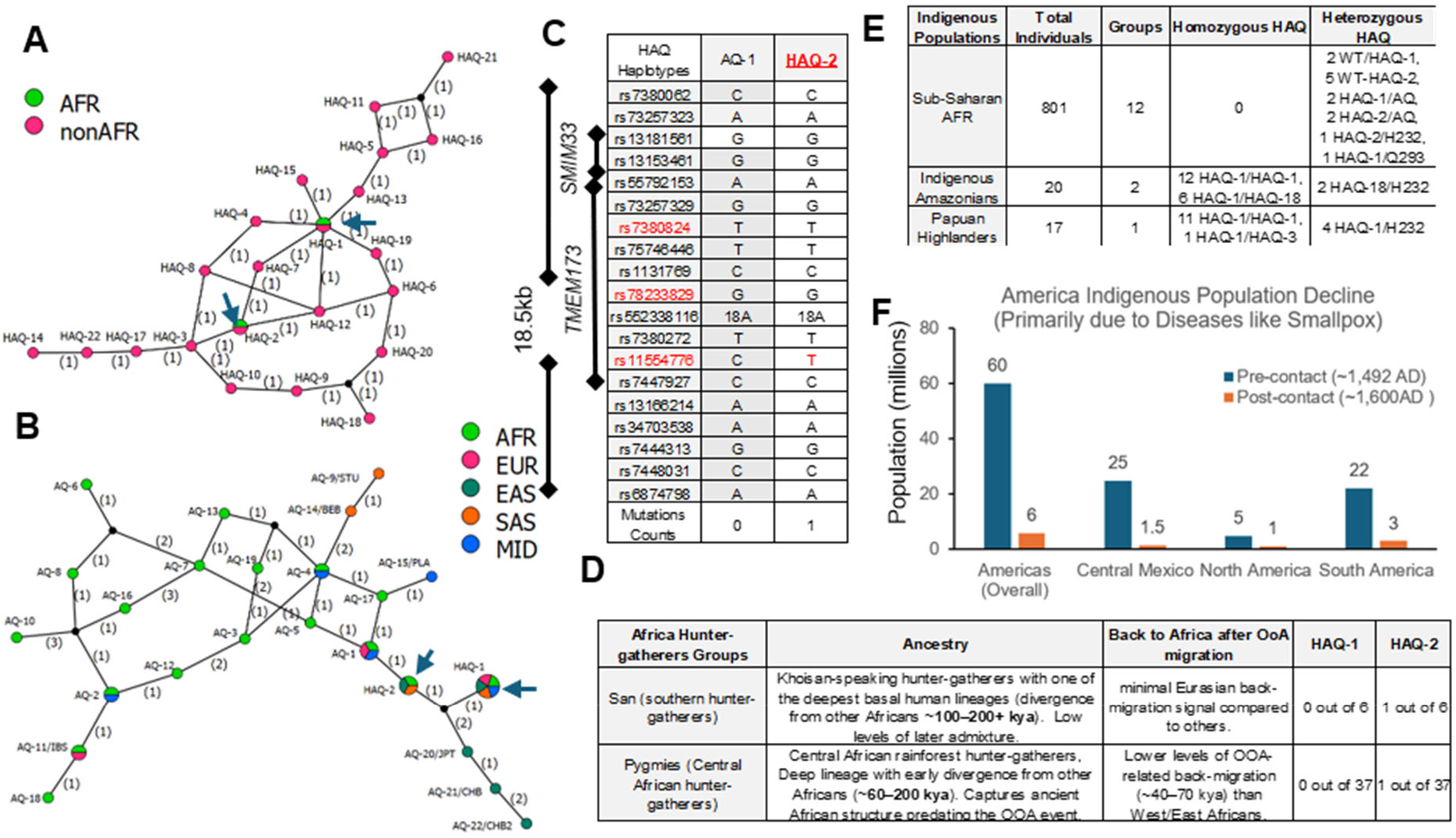
The *HAQ* allele is negatively selected in Sub-Saharan Africa. **A-B**. A network of *HAQ* (all 22 haplotypes) and *AQ* (all 22 haplotypes plus the *HAQ-1, HAQ-2* haplotypes) found in 1KGP+HGDP individuals was constructed using Popart (popart.otago.ac.nz) by TCS network method (*49*). The number on the line indicates the number of mutations separating the adjacent haplotypes. **C**. Comparing the 18.5 kb *STING* haplotype consisting of 19 SNPs in *AQ-1, HAQ-2*. **D**. Examine *HAQ-1, HAQ-2* haplotypes in African hunter-gatherer groups, the San and Pygmies. **E**. Compare *HAQ* genotypes in indigenous groups living in rainforests: Sub-Saharan Africans, Amazon indigenous groups, and Papuan highlanders. **F**. Compare native American populations pre- and post-Columbus era, showing population decrease due to the smallpox epidemic.

### The positively selected *HAQ-1* haplotype evolved at the Southern Indian Coast at least 50,000 years ago

The *HAQ-1* haplotype is by far the dominant *HAQ* haplotype in all non-Africans (*11*) (Figure S1). The high population frequency and long linkage disequilibrium (18.5kb) consistently signal positive natural selection of *HAQ-1* during the OoA migration. We then investigate the evolution of the *HAQ-1* haplotype. Papuan highlanders carry the *HAQ-1* and *HAQ-3* haplotypes (Figure 2E). We reason that *HAQ-1* and *HAQ-3* arose before the separation of Papuan highlanders and EAS branches during OoA migration, some 50kya ago (*33-37*) (Figure 3A). AFR has the *HAQ-1* and *HAQ-2* haplotypes, but not *HAQ-3. HAQ-1* is two mutations away from *HAQ-2* and also cannot be derived from any *AQ* haplotype in Africa (Figure 2B). The transitional *HAQ* haplotypes from *HAQ-2* to *HAQ-1*, i.e. *HAQ-7* and *HAQ-12*, only exist outside Africa (Figure 2A, 3B). In contrast, the *HAQ-3* haplotype is 1 mutation away from *HAQ-2* (Figure 3C). Lastly, the population frequency of *HAQ-1* in AFR is only 0.25% (Figure S1). We reasoned that *HAQ-1* evolved from *HAQ-2* outside Africa. *HAQ-1* in AFR arose from back migration to Africa during the OoA migration.

**Figure 3.**
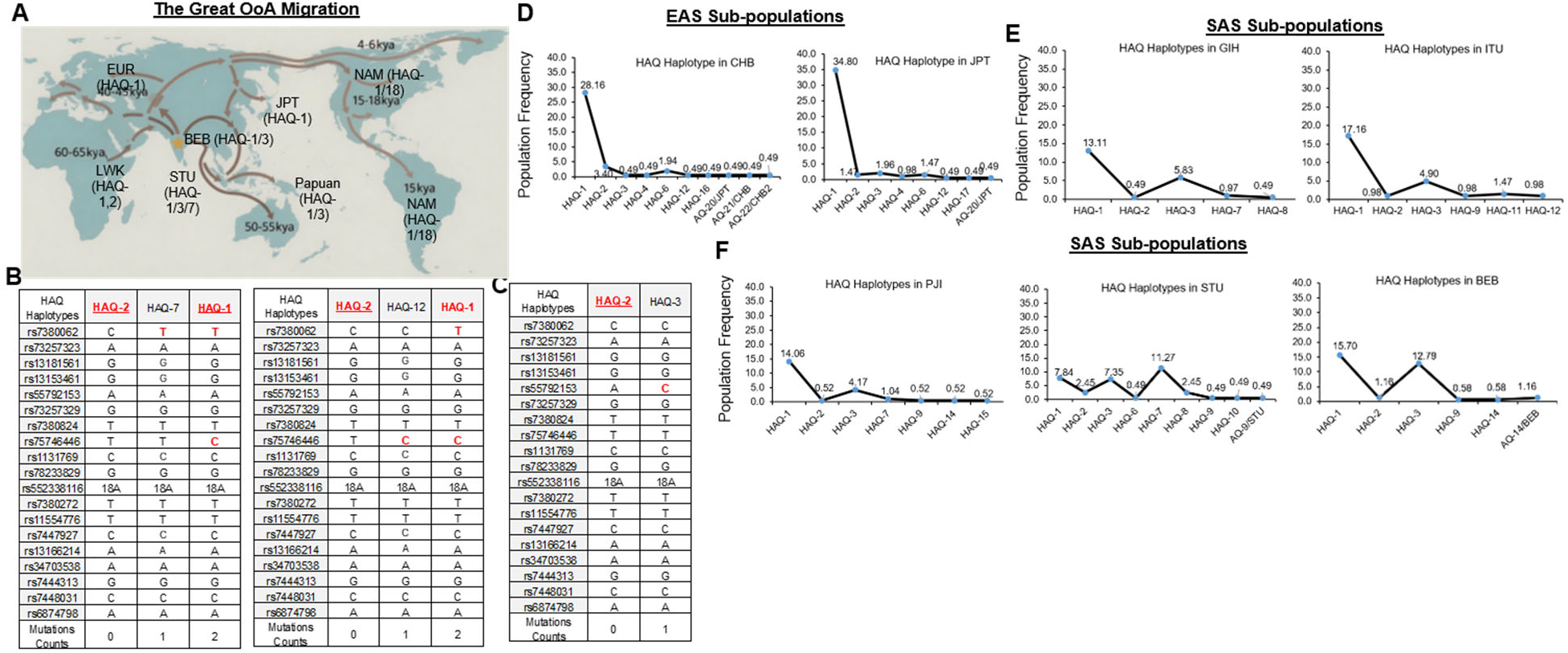
The *HAQ-1* haplotype evolved on the Southern Indian Coast 50,000 years ago. **A**. The southern coast route of OoA migration. The arrow indicates the route. The possible branch deviation time is indicated. Image is modified from Genome Research Limited – Wellcome Sanger Institute. STU: Tamil from Sri Lanka; BEB: Bengali in Bangladesh. **B-C**. Comparing the 18.5 kb *STING* haplotype consisting of 19 SNPs in *HAQ-1, HAQ-2*, and *HAQ-3*. **D-F**. The population frequency and genetic diversity of the 18.5 kb *STING* haplotype were analyzed in *HAQ* individuals using 19 SNPs in South Asians (SAS), and East Asians (EAS) in 1kgp+HGDP groups.

AMH left Africa around 50,000–70,000 years ago via the Bab-el-Mandeb strait or nearby coastal areas, then following the southern coastline of Asia eastward (*39, 40*) (Figure 3A). The Oceania (Australo-Papuan branch) is the earliest split. Ancestors of Aboriginal Australians and Papuans diverged from other Eurasians ~50,000–72,000 ya (*33-37*). East Asian branch diverged from West Eurasians around 50,000–40,000 ya (Figure 3A). Central Asian branch, ancestral to some North Asian and Native American lineages, diverged ~45,000–30,000 ya. The West Eurasian branch moved northward/westward into the Near East and Europe ~55,000–45,000 ya (*41-43*) (Figure 3A). Since *HAQ-1* existed before 50,000 ya, it must have evolved along the southern Indian coast during the OoA migration before the EAS-OCE or East-West Eurasian split. Our *HAQ* haplotype analysis (Figure 3E) also supports this conclusion.

We examined *HAQ* haplotypes in all 62 non-African groups. *HAQ-1* was clearly the #1 *HAQ* haplotype in 60 of the 62 groups (Figure S1, Figure 3D), supporting the notion that *HAQ-1* evolved early and served as the founder for most non-African groups. The two exceptions are the STU group (Tamil from Sri Lanka, sampled in the UK) and the BEB group (Bengali in Bangladesh) (Figure 3E, 3F). STU has three dominant *HAQ* haplotypes: *HAQ-1* (7.84%), *HAQ-3* (7.35%), and *HAQ-7* (11.27%) (Figure 3F). The BEB group has two dominant *HAQ* groups: *HAQ-1* (15.7%) and *HAQ-3* (12.79%) (Figure 3F). Thus, unlike the other non-African groups, the STU and BEB groups had multiple *HAQ* founders: *HAQ-1, HAQ-3*, and *HAQ-7*. We concluded that these *HAQ* founders evolved on the southern Indian coast near modern-day Sri Lanka and Bangladesh. Moreover, the fact that *HAQ-1, HAQ-3, and HAQ-7* were similarly selected on the southern Indian coast indicated that various *HAQ* haplotypes are functionally comparable.

### The positively selected *HAQ* had more human generations than the neutral *H232* during the OoA migration

How did the *HAQ* allele rise from ~1% to ~80% in NAM during OoA migration? We compared the evolution of the positively selected *HAQ* haplotypes with that of the neutral *H232* haplotypes during OoA migration (Figure S2A, S2B). The AFR has 4 *H232* haplotypes, with *H232-1* as the dominant haplotype (Table S3, Figure 4A). In non-Africans, an additional 8 *H232* haplotypes show a pattern of classic genetic drift (Figure 4B). *H232-1* remains the #1 haplotype across the groups, indicating that *H232-1* was the founder outside Africa (Figure 4A, 4B). With founders identified, we next drew phylogenetic trees to compare the evolution of *H232* and *HAQ* outside of Africa.

**Figure 4.**
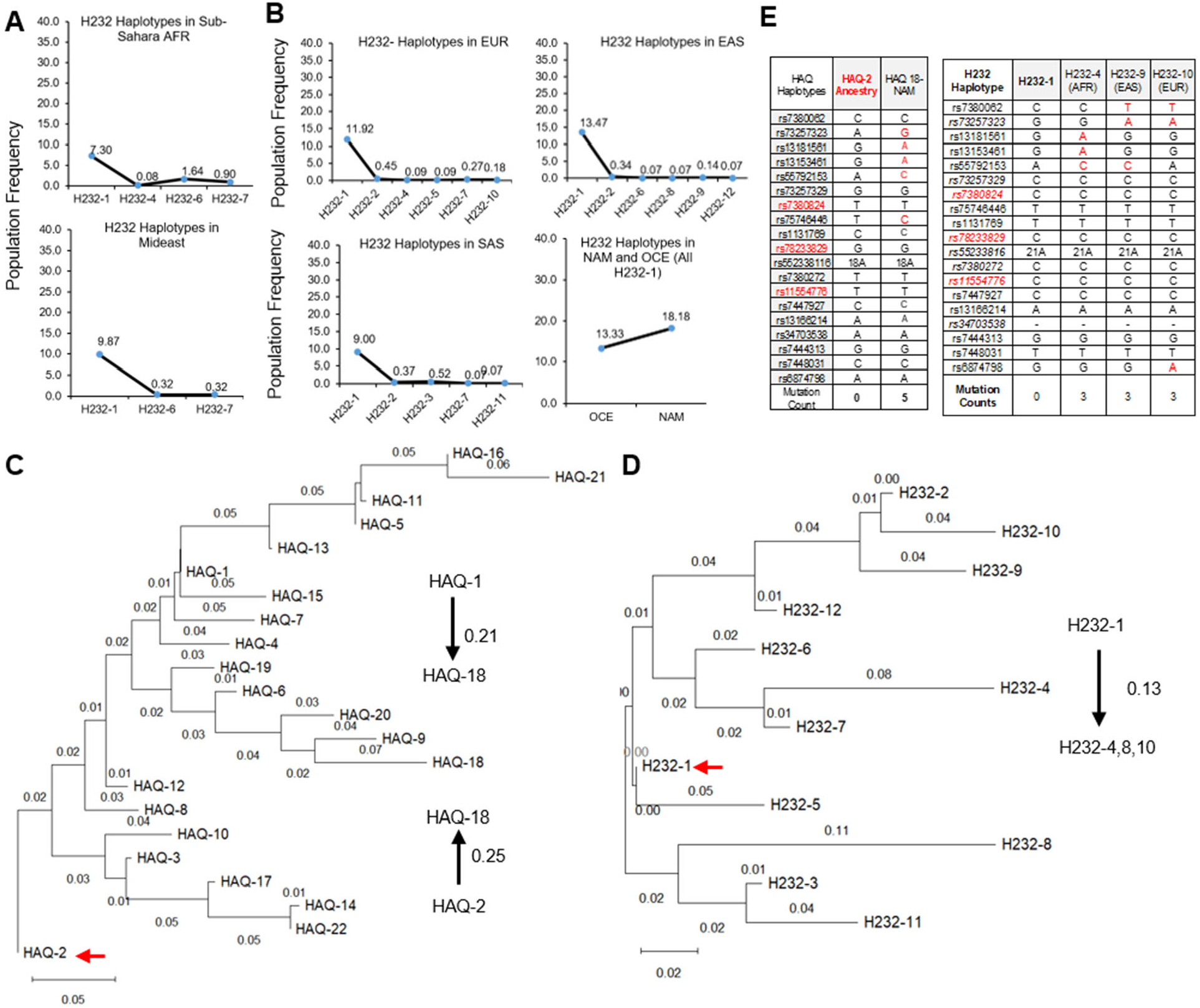
*The HAQ carriers had more generations than the H232 carriers during the out-of-Africa migration*. **A-B**. The population frequency and genetic diversity of the 18.5 kb STING haplotype were analyzed in *H232* individuals using 19 SNPs across 1kgp+HGDP groups, including Sub-Saharan Africa, the Middle East, Europe, South Asians, East Asians, and Native Americans. **C-D**. Phylogenetic analysis of all 22 HAQ and 12 H232 haplotypes was performed using MEGA 12 software. The evolutionary history was inferred using the Neighbor-Joining method (50). The percentage of replicate trees in which the associated taxa clustered together in the bootstrap test (500 replicates) is shown next to the branches (51). The tree is drawn to scale, with branch lengths (next to the branches) in the same units as those of the evolutionary distances used to infer the phylogenetic tree. The evolutionary distances were computed using the p-distance method (52) and are in the units of the number of base differences per site. The analytical procedure encompassed 22 nucleotide sequences. The pairwise deletion option was applied to all ambiguous positions for each sequence pair, resulting in a final data set comprising 19 positions. Evolutionary analyses were conducted in MEGA12 utilizing up to 8 parallel computing threads. Founder haplotypes, *HAQ-2* and *H232-1*, are indicated by the red arrow. The branch distance is shown.

In the *HAQ* tree, *HAQ-2* is the founder. The three long branches are *HAQ-2* to *HAQ-14* (branch length 0.17), *HAQ-2-HAQ-18* (0.25), and *HAQ-2-HAQ-1-HAQ-21* (0.31). In the *H232* tree, *H232-1* is the founder. The three long branches are *H232-1* to *H232-11* (0.13), *H232-1* to *H232-4* (0.13), and *H232-1* to *H232-10* (0.14). The *HAQ* phylogenetic tree is longer than the *H232* tree in non-Africans (Figure 4C, 4D). The PopArt maps also showed that *HAQ* haplotypes are more distal than *H232* haplotypes (Figure S2A, S2B).

To estimate the molecular clock, we examined accumulated mutations across distal haplotypes. For the *HAQ* tree, we selected the NAM-specific *HAQ-18* haplotype as the distal *HAQ* haplotype. *HAQ-18* is carried by 18.55% of NAM (Figure S1B), 12.5% of Mexicans, and 16.47% of Peruvians (Figure S2C). In fact, *HAQ-18* is the 2^nd^ most common *HAQ* haplotype next to *HAQ-1* in all groups (Figure S1B). Thus, *HAQ-18* represents a distal but relevant *HAQ* haplotype. We counted 5 mutations from *HAQ-2* to *HAQ-18* (Figure 4E) or from *HAQ-1* to *HAQ-18* (Figure S2D). In contrast, we counted 3 mutations from *H232-1* founder to the three distal *H232* haplotypes, *H232-4, H232-8*, or *H232-10* (Figure 4E), Thus, the *HAQ* allele accumulated more mutations than the *H232* allele in the same 18.5kb *STING* genomic region during OoA migration. The accumulated mutations= *numbers of mutations per generation x (total time outside Afiica*)*/*(*generation time*).

The *HAQ* and *H232* alleles have the same mutation rate since they are in the same *STING* gene region. The total time is also the same for the *HAQ* and *H232* alleles outside Africa. We concluded that *HAQ* allele carriers had a shorter generation time than the *H232* carriers during the OoA migration.

To further demonstrate that *HAQ* carriers had a short generation time during the OoA migration, we examined generation time in the *WT* carriers. There are 33 *WT* haplotypes in 1KGP+HGDP (Table S4). 18 are in AFR (Figure S3A), indicating that *WT* haplotypes in non-Africans had multiple founders. We could not draw a *WT* phylogenetic tree for non-Africans. Nevertheless, across all 33 WT haplotypes, we found that all non-African haplotypes are within 3 mutations of an AFR WT haplotype (Figure S3B). For example, the distal non-AFR *WT-18* haplotype is two mutations away from the AFR *WT-17* haplotype (Figure S3C). Similarly, the distal non-AFR *WT-30* haplotype is three mutations away from the AFR *WT-12* haplotype (Figure S3C). Thus, during OoA migration, the *WT* allele accumulated a similar number of mutations to the *H232* allele. The *HAQ* allele evolved faster than either the *H232* or the *WT STING* allele during OoA.

### Forward simulation based on the molecular clock difference between *HAQ* and *H232* predicts elevated population *HAQ* frequency during OoA migration

All the mutations in the *HAQ* and *H232* haplotypes are in the introns. The excess of intronic mutations on *HAQ* is quantitatively consistent with its elevated time-integrated frequency under positive selection after the OoA dispersal. To link the molecular signature (five intronic mutations on *HAQ* versus three on *H232* in the same genomic region and over the same period during OoA) to population genetic patterns, we performed forward simulations using the mutation difference as the primary independent constraint. *H232* was modeled as neutral starting at ~20% frequency. *HAQ* started at 2%. The selection coefficient was tuned so that the cumulative time-integrated frequency (∑ p(t), total haplotype-generations) of *HAQ* was ~1.67× higher than that of *H232*, directly reflecting the observed 5:3 mutation ratio. The simulation showed that *H232* (blue dot line) is neutral and remains stable near 20%. *HAQ* (red line) starts at 2% under positive selection with s = 0.00185 (Figure 5A). The selection coefficient was adjusted solely to satisfy the observed intronic mutation accumulation ratio

**Figure 5.**
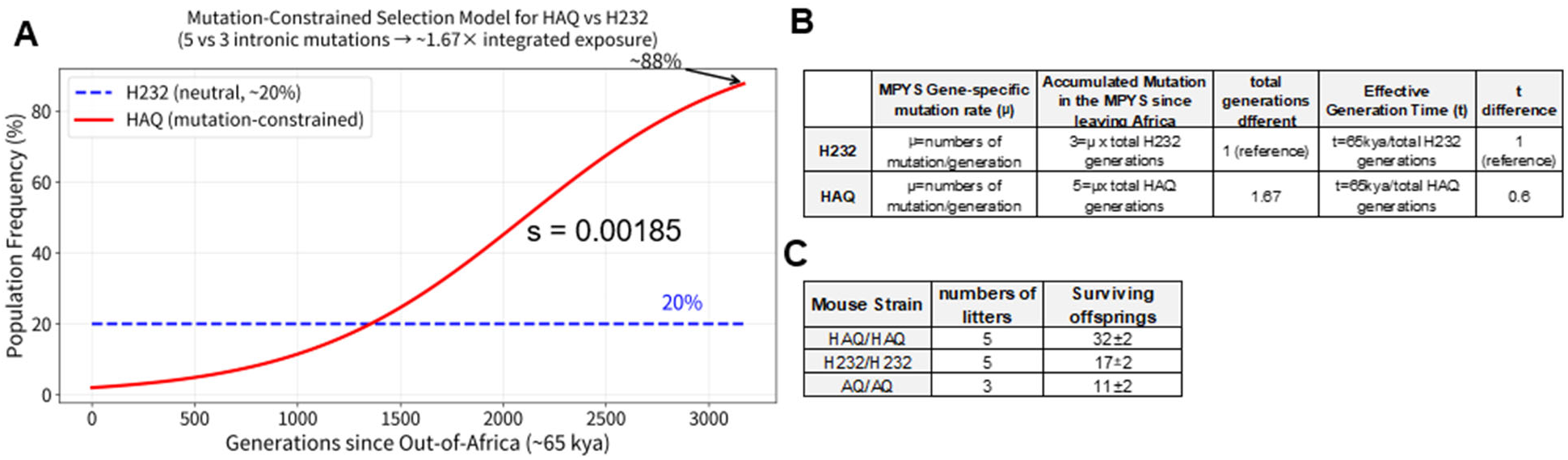
Forward simulation and mouse models illustrate the short effective generation time in *HAQ*, leading to positive selection in OoA migration. **A**. Mutation-constrained selection model for *HAQ* vs *H232* allele frequency trajectories since OoA migration (Material and Methods). *H232* (blue dashed line) is modeled as neutral and remains stable near 20%. *HAQ* (solid red line) starts at 2% under positive selection. The selection coefficient (s=0.00185) was tuned solely to satisfy the observed intronic mutation accumulation ratio (5 mutations on *HAQ* vs. 3 on *H232*), corresponding to ~1.67× higher cumulative time-integrated frequency for *HAQ*. This mutation-driven model independently predicts a strong rise in *HAQ* frequency, reaching ~88% after ~3,170 generations (~65 kya at 20.5 years/generation). **B**. A method to calculate the generational time difference between *HAQ* and *H232* during OoA migration. **C**. Breeding record of the *HAQ, H232, AQ* mice in a SPF facility over a 6-month period.

This mutation-driven model predicts a substantial increase in *HAQ* frequency reaching 88% over ~3,170 generations (~65 kya at 20.5 years/generation (*29*)), consistent with the observed modern cline (*H232* stable at ~17–27% across non-African populations; *HAQ* reaching ~82% in NAM). Although the deterministic model provides a simplified representation, including realistic stochastic drift and serial founder effects during OoA dispersal and subsequent migrations would be expected to produce additional variance while preserving the overall upward trajectory of *HAQ*. The close correspondence between the mutation-constrained prediction and empirical frequencies supports the conclusion that positive selection on *HAQ* drove its rise in non-African populations, with the excess intronic mutations serving as a molecular record of *HAQ*’s shorter generation time.

### The effective generation time for *HAQ* carriers was 60% of that for *H232* carriers during the OoA migration

Using the number of mutations as a molecular clock, we calculated the generation time difference between *HAQ* and *H232* carriers. The mutation rate is the same for the two alleles because they are in the same *STING* genomic region. The total time passed is the same, i.e. the time of OoA migration. During OoA migration, the generation time for *HAQ* carriers was 60% of that for *H232* carriers (Figure 5B). Notably, this is the effective generation time, from birth to the point at which they produce surviving offspring. It was estimated that AMH had an average generation time of roughly 27 years over the past ~250,000 years (*29*), but this varied significantly by population, sex, and time period. Obviously, AMH can have offspring much earlier than 27 years of age, indicating that the generation time in early AMH was constrained by other factors, such as nutrition and disease, that determine offspring survival. To examine these variables influencing the effective generation time, we determined reproduction using mouse models of *HAQ* and *H232*.

### *HAQ* mice produce more surviving offspring than *H232* mice in an SPF mouse facility

*HAQ* and *H232* knock-in mice were previously generated by us (*10, 44*). We bred 8-week-old *HAQ/HAQ* or *H232/H232* mice at a 1:1 male-to-female ratio for 6 months. The mice were housed in the same special pathogen-free (SPF) room, under the same 12hr light-dark cycle. All mice were fed the Teklad Irradiated Global 18% Protein, 6% fat Rodent Diet (Inotiv, cat# 2918). We recorded the total number of litters and surviving pups. Both strains produced 5 litters on average during the 6-month period. However, the *HAQ/HAQ* generated an average of 32 surviving pups compared to 17 pups by *H232/H232* (Figure 5C). Thus, the *HAQ* mice produce more surviving offspring than the *H232* mice, consistent with the positive selection of *HAQ* in OoA migration. Moreover, *HAQ* was positively selected in the absence of disease. We also recorded *AQ/AQ* breeding for 6 months. The *AQ* allele was negatively selected during OoA migration (*11*). The *AQ/AQ* colony had only 3 litters and, on average, 11 surviving pups over a 6-month period (Figure 5C).

## Discussion

The HAQ protein has three amino acids difference from the WT STING protein: R71H-G230A-R293Q (*9*). In comparison, the WT human and chimpanzee STING proteins differ by only two amino acids: R78W and G230A (*11*). The human and chimpanzee share a common ancestry 6 ~7 million years ago. Here, we describe the opposing selection of *HAQ STING*: purifying selection in Sub-Saharan Africans and positive selection in non-Africans.

What causes the purifying selection in Sub-Saharan Africans but is absent in NAM? We initially described the *HAQ* as a loss-of-function *STING* allele because it is defective in the production of type I IFNs (*9, 10*). Notably, PBMCs from smallpox-vaccinated *HAQ* humans are defective in the IFNα response to poxviruses (*45*). Thus, smallpox is particularly lethal to the *HAQ* individuals due to the lack of type I IFNs production. Smallpox was the #1 cause of population collapse in NAM post-Columbus. The indigenous American population decreased from 60 million to 6 million (Figure 2F). Here, we found that 51.61% of NAM are *HAQ/HAQ*, compared with 2.23% in EUR and 0% in AFR, making NAM highly susceptible to smallpox and possibly other infectious diseases (e.g., measles), which together killed millions in NAM. Moreover, today’s NAM are descendants of the epidemic survivors. Thus, pre-1492 *HAQ/HAQ* frequency might have been higher. Indeed, in two isolated indigenous Amazon groups, Surui and Karitiana, the *HAQ/HAQ* carriers reach ~90% (Figure 2E).

Smallpox (caused by the variola virus) originated from a zoonotic spillover of an African rodent poxvirus (*46*). Sub-Saharan Africans have evolved alongside endemic disease caused by zoonotic spillover, which likely is the purifying force that negatively selected *HAQ/HAQ* in Sub-Saharan Africans. Furthermore, the purifying pathogen must have high mortality in AMH, i.e., a highly virulent strain. Viruses that establish latency in AMH, as in the Herpesviruses EBV, CMV, and HSV, are unlikely to be the purifying force for *HAQ* individuals. The most virulent human virus strains often jump directly from animal hosts to humans. Considering early humans were hunter-gatherers in small groups, we reason that viruses that negatively selected *HAQ* in Sub-Saharan Africans had an animal reservoir. When early AMH left Africa ~70,000ya, they were separated from these animal reservoirs. Thus, the purifying force on *HAQ* was lifted.

*HAQ STING* shows a modest, continuous positive selection during the OoA migration. Which *STING* function was positively selected in non-Africans? Unlike the purifying selection observed in Africa, it is not related to pathogen infection, since the HAQ mouse housed in an SPF environment still produced more surviving offspring. How did *HAQ* generate more surviving offspring? Maternal and infant nutrition during pregnancy, lactation, and the first two years of life is one of the most critical factors influencing a child’s survival, growth, and long-term health. Undernutrition is a major contributor to child mortality, responsible for about 45% of deaths in children under 5 worldwide today. Breastfeeding, in particular, is a key protective factor providing essential nutrients, bioactive compounds, and immune protection that lower infant mortality and improve cognitive development. Fatty acids are a major component of breast milk, making up approximately 3% to 5% of its weight and providing around 50% of the energy required by infants in the first months of life. Essential fatty acids like linoleic acid (omega-6) and alpha linolenic acid (omega-3) cannot be synthesized by infants and must come from breast milk. We reported that HAQ mice have more fat storage and regulate saturated vs unsaturated fatty acid production (*11, 47*). WT STING can inhibit fatty acid desaturase 2 (FADS2), the rate-limiting enzyme in polyunsaturated fatty acid (PUFA) desaturation (*47*). PUFA synthesis deficiency caused infertility in fads2^-/-^ female mice (*48*). Sustained supply of PUFA restored fertility in adult infertile fads2^-/-^ mice (*48*). The breeding defect is more severe in the AQ mice, which had fewer litters. In summary, good maternal and infant nutrition is a cornerstone of infant survival. We hypothesize that *HAQ* increases infant survival by enhancing fatty acid species and amounts in breast milk.

The selection coefficient shows regional differences: highest in NAM (0.00178), lowest in EUR (0.00099). We attribute this to a balance of negative and positive selection in these regions. The American continent had no purifying selection; thus, positive selection for *HAQ/HAQ* was in full force, driving *HAQ/HAQ* to over 50%. Notably, in Papuan highlanders and indigenous Amazon groups, *HAQ/HAQ* reaches 70%~90% and even eliminates the *WT STING* genotype. The West Eurasian branch of OoA migration separated about the same time as EAS and Papuan, around 45~50kya ago (*41-43*). In contrast, *HAQ/HAQ* only accounts for 2% of Europeans. From the outcome of the 1492 Columbus encounter, we can confirm that *HAQ* faced a purifying force in Europe after the OoA migration. The low S (0.00099) in Europeans thus reflects the combined effect of positive and negative selection.

What does the natural selection of *HAQ* during the OoA migration mean for AMH today and in the future? Vaccines, antibiotics, clean water, food safety, and personal hygiene have eliminated the purifying force for *HAQ* in most societies. Smallpox was eradicated in 1980. But in places where zoonotic pathogens are still prevalent, for example, Sub-Saharan Africa, *HAQ/HAQ* individuals are still at high risk. Modern healthcare has also greatly improved infant survival. Positive selection for *HAQ*, with more surviving offspring, is likely diminished, though India and China, which have a majority of *HAQ* carriers, remain the two most populous countries, with 2.8 billion out of a total of 8.3 billion AMH today. Lastly, we observed that *HAQ/HAQ* became fixed in isolated Papuan highlanders and indigenous Amazon groups. Very few *WT/WT* exist in these isolated groups. Notably, the Papuans developed their own agriculture. Thus, it was not just the hunter-gatherers in the OoA migration. It seems that, in the absence of purifying selection, positive selection will drive *HAQ* to fixation in AMH, becoming the new *WT*. “*What’s past is prologue*” (The Tempest). Natural selection made us who we are today and will continue to shape who we will become in the future.

### The limitations of the study

The purifying and positive selection are supported by studies in 74 groups, ~4,000 AMH. The molecular clock difference in OoA supported the difference in generation times. The mechanism behind the generation time difference was based on a mouse study under SPF conditions. No causal relationship has been established between increased pup survival and enhanced fatty acid metabolism in *HAQ*.

## Acknowledgments

Grok (xAI) provided guidance on Hail MatrixTable queries, Dataproc cluster setup, and Python scripting for genotype extraction and allele conversion.

## Author contributions

Conceptualization: LJ

Methodology: LJ

Investigation: AAT, LJ

Supervision: LJ

Writing—original draft: LJ

Writing—review & editing: AAT, LJ

## Data, code, and materials availability

All data are available in the main text or the supplementary materials.”

## Supplementary Materials

Materials and Methods

Figs. S1 to S3

Tables S1 to S4

