## Supplemental File for "Natural Selection of the Common *STING* Allele *HAQ* in Anatomically Modern Humans"

**The PDF file includes:**

Materials and Methods
Figs. S1 to S3
Tables S1 to S4

### Materials and Methods

**Experimental Design.** The study was designed to answer how the *HAQ* allele was naturally selected during OoA migration. To cover the evolution of the *HAQ* allele, we include data from HGDP, which contains key groups during OoA, e.g., the Mideast, Papuan highlander, and Native Americans, that are missing from the 1KGP. The study uses individual-level genotype data, which are essential for estimating the molecular clock of *HAQ* evolution. Computer simulation was used to confirm the positive selection of *HAQ* during OoA migration. A knock-in mouse model of *HAQ* was used to confirm the difference in generation time. Mouse experiments were biological replications that involved the same experimental procedures on different mice.

### Methods

**Genotype Extraction.** Individual-level genotype data for 19 target SNPs on chromosome 5 were extracted from the gnomAD v3.1.2 genomes dataset using the Hail 0.2.138 framework on Google Cloud Dataproc. The public dense MatrixTable (gnomad.genomes.v3.1.2.hgdp_1kg_subset_dense.mt) was accessed directly from Google Cloud Storage without local downloading of large VCF files. A Dataproc cluster (master node: n1-highmem-8, 2 worker nodes: n1-standard-8) was provisioned in the us-central1 region using hailctl dataproc start. The Compute Engine default service account was granted necessary IAM roles (Dataproc Editor/Worker, Storage Object Viewer/Creator, Logs Writer, Monitoring Metric Writer, and Service Account User) to enable cluster creation and GCS access. For each SNP, the MatrixTable was filtered to HGDP samples only (s.startswith('HGDP')), then to the specific genomic position using hl.locus ('chr5', position, reference_genome='GRCh38'). Genotype entries were extracted as a Hail Table (entries().key_by('s').select('GT')) and written to a Hail Table (.ht) format. Genotypes were converted from numeric/phased format (0/0, 0/1, 1/1) to real allele notation (e.g., C/C, C/T) using custom Python mapping based on the reference (REF) and alternate (ALT) alleles from gnomAD. Final tables were exported as CSV files using pandas for downstream analysis. This pipeline allowed efficient extraction of per-sample genotypes from the full HGDP cohort (~828 samples) without MAF filtering or large local storage requirements. All scripts were developed iteratively in the Hail Python environment on the Dataproc master node. Individual-level genotype data from 1kgp was downloaded as before [11].

**Estimation of selection coefficients and confidence intervals.** Allele frequencies for HAQ were extracted directly from the population frequency in 1KGP+HGDP. Frequencies were recorded for the following population groups: Sub-Saharan African (AFR), Non-Finnish European (NFE), Finnish (FIN), Middle Eastern (MID), South Asian (SAS), East Asian (EAS), and Native American (NAM). Selection coefficients (s) were estimated under a deterministic logistic (genic/additive) selection model. For each target population, the per-generation selection coefficient was calculated as:

$$s \approx \frac{\ln\left(\frac{p_1}{1-p_1}\right) - \ln\left(\frac{p_0}{1-p_0}\right)}{t}$$

where $p_0$ is the ancestral frequency (Sub-Saharan African value, 1.62%), $p_1$ is the observed frequency in the target population, and t is the number of generations since the Out-of-Africa dispersal. We used a consensus divergence time of approximately 65,000 years before present and an ancestral generation time of 20.5 years for non-African lineages, yielding t ≈ 3,171 generations. Branch-specific estimates incorporated published split times: West Eurasian–East Eurasian split ≈ 45 kya and East Asian–Native American split ≈ 22 kya. Approximate 95%

confidence intervals for s were obtained via Monte Carlo simulation (10,000 iterations). In each iteration, $p_0$ was drawn from Normal (0.0162, 0.005), modern frequencies $p_1$ incorporated minor sampling variance, and total generations t were drawn from Normal (3,171, 400) to reflect uncertainty in Out-of-Africa timing (60–70 kya) and generation interval (20.5 ± 2–3 years). The 2.5th and 97.5th percentiles of the resulting s distribution were reported as the approximate 95% CI. All calculations were performed in Python 3 using NumPy. Fold increases, odds ratios, and summary statistics for *HAQ* (mean ± SD across non-African groups) were computed directly from the extracted frequencies. The logistic trajectory was simulated by iterating the recurrence $p_{t+1}$ = $p_t \times (1+s)$ / $1 + s \times p_t$ to confirm consistency with observed clines.

**Forward simulation of allele frequency trajectories.** To link the observed difference in intronic mutation accumulation (5 mutations on HAQ versus 3 on H232) to population frequency changes, we performed deterministic forward simulations over 3,170 generations, corresponding to ~65,000 years at an ancestral non-African generation time of 20.5 years per generation [37].H232 was modeled as neutral with frequency held constant at its starting value of ~20%. HAQ was modeled as starting at 2% frequency under positive selection using the logistic (genic) selection recurrence relation:

$$p_{t+1} = \frac{p_t \times (1+s)}{1 + s \times p_t}$$

To avoid circularity with modern frequency observations, the primary modeling constraint was the observed difference in accumulated intronic mutations (5 intronic mutations on *HAQ* versus 3 on *H232*). Because the expected accumulation of neutral mutations on a haplotype class is proportional to its time-integrated frequency (∑ p(t) across generations), the single selection coefficient s was iteratively adjusted until the cumulative haplotype-generations experienced by *HAQ* was ~1.67× higher than that of *H232*. The final value used was s = 0.00185 per generation. Cumulative exposure for each allele was calculated as the sum of its frequency p(t) over all 3,170 generations. The resulting trajectories were plotted to visualize the predicted change in population frequency since the OoA dispersal.

Python code used for the simulation:

```
python
import numpy as np
import matplotlib.pyplot as plt
# Parameters
generations = 3170
s = 0.00185          # mutation-constrained selection coefficient
p_H232 = 0.20        # neutral, stable frequency
p_HAQ = 0.02         # starting frequency for HAQ
# Arrays to store frequencies
freq_H232 = np.zeros(generations + 1)
```

```python
freq_HAQ = np.zeros(generations + 1)
freq_H232[0] = p_H232
freq_HAQ[0] = p_HAQ
# Simulate forward
for t in range(generations):
    freq_H232[t+1] = freq_H232[t]  # neutral
    # Logistic selection for HAQ
    p = freq_HAQ[t]
    freq_HAQ[t+1] = p * (1 + s) / (1 + s * p)
# Calculate time-integrated exposure (key constraint)
integrated_H232 = np.sum(freq_H232)
integrated_HAQ = np.sum(freq_HAQ)
ratio = integrated_HAQ / integrated_H232
print(f"Time-integrated ratio (HAQ/H232): {ratio:.3f}x")
print(f"Final HAQ frequency: {freq_HAQ[-1]*100:.1f}%")
# Plotting
plt.figure(figsize=(10, 6))
plt.plot(freq_H232 * 100, color='blue', linestyle='--', label='H232 (neutral, ~20%)')
plt.plot(freq_HAQ * 100, color='red', label='HAQ (mutation-constrained)')
plt.axhline(y=20, color='blue', linestyle=':', alpha=0.5)
plt.xlabel('Generations since Out-of-Africa (~65 kya)')
plt.ylabel('Population Frequency (%)')
plt.title('Mutation-Constrained Selection Model for HAQ vs H232\n'
          '(5 vs 3 intronic mutations → ~1.67× integrated exposure)')
plt.legend()
plt.grid(True, alpha=0.3)
# Annotate final frequencies
plt.annotate(f'~88%', xy=(generations, freq_HAQ[-1]*100),
             xytext=(generations-100, freq_HAQ[-1]*100 + 3),
             arrowprops=dict(arrowstyle='->', color='black'), fontsize=12,
color='black')
plt.annotate('20%', xy=(generations, 20),
             xytext=(generations-200, 23), fontsize=12, color='blue')
```

```
plt.tight_layout()
plt.show()
```

**Mice**. *HAQ*, *H232*, *AQ* mice were previously generated in the lab [10,11,48]. Mouse breeding includes one male to one femal (2- month old), >4 pairs for every breeding over 6 months period. Mice were housed at 22°C under a 12-h light-dark cycle with ad libitum access to water and a chow diet (3.1 kcal/g, Teklad 2018, Envigo, Sommerset, NJ) and under pathogen-free conditions in the Animal Research Facility at the University of Florida. Litter numbers and surviving litter were record. All mouse experiments were performed by the regulations and approval of the Institutional Animal Care and Use Committee at the University of Florida, IACUC202200000058.

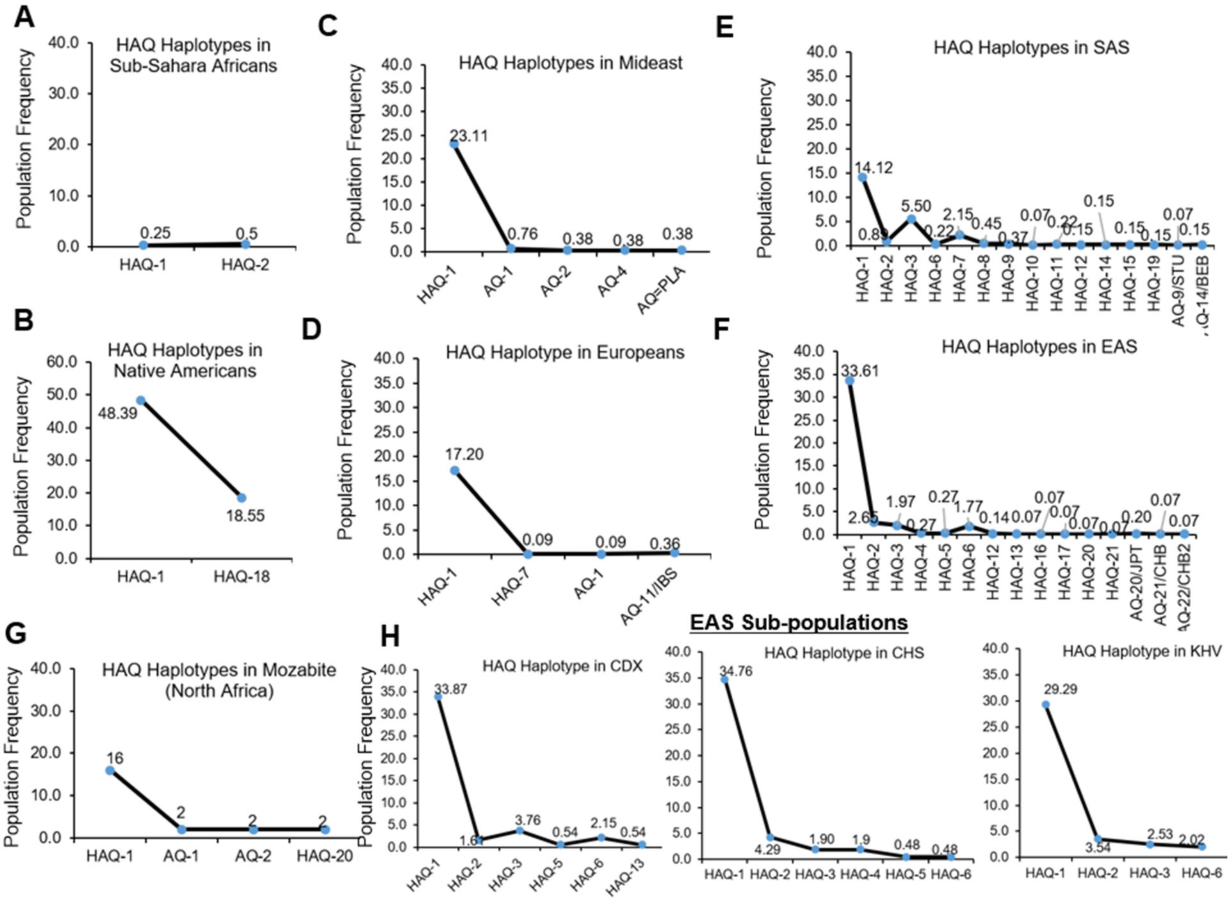

**Figure S1.** ***HAQ-1*** **is the dominant** ***HAQ*** **haplotype, followed by the NAM-specific** ***HAQ-18*****.** **A-H.** The population frequency and genetic diversity of the 18.5 kb *STING* haplotype were analyzed in *HAQ* individuals using 19 SNPs across 1kgp+HGDP groups, including Sub-Saharan Africa, the Middle East, North Africans, Europeans, South Asians, East Asians, and Native Americans.

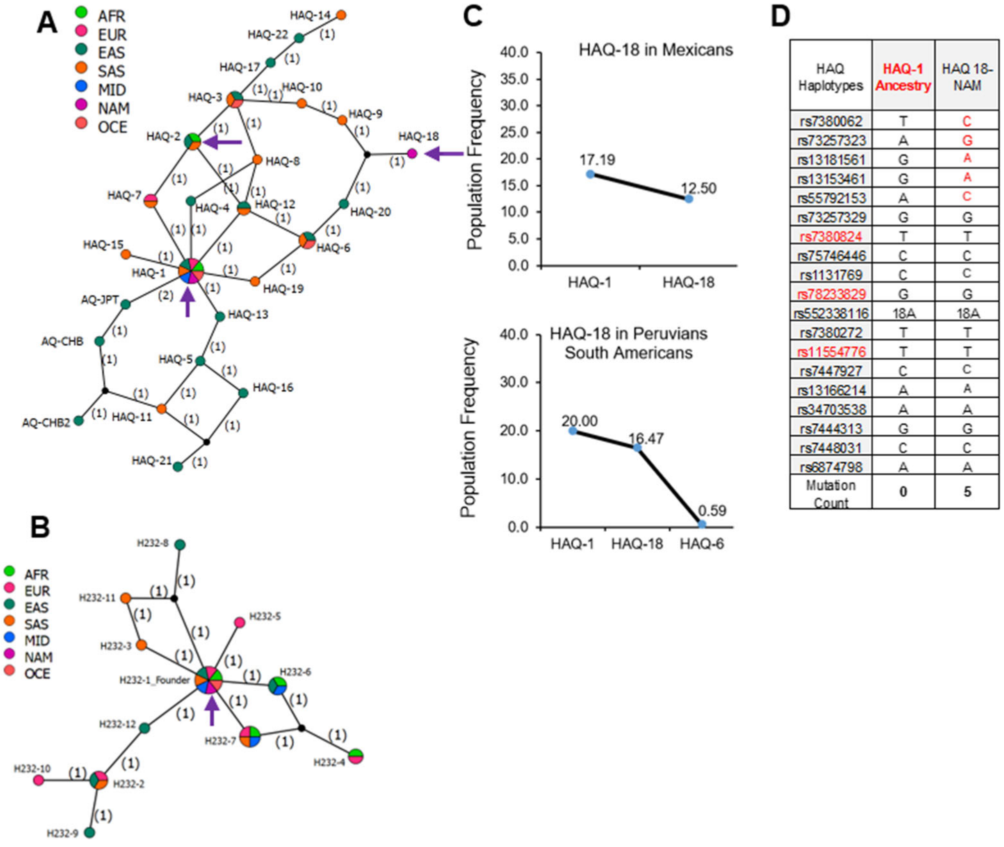


| HAQ Haplotypes | HAQ-1 Ancestry | HAQ 18-NAM |
|---|---|---|
| rs7380062 | T | C |
| rs73257323 | A | G |
| rs13181561 | G | A |
| rs13153461 | G | A |
| rs55792153 | A | C |
| rs73257329 | G | G |
| rs7380824 | T | T |
| rs75746446 | C | C |
| rs1131769 | C | C |
| rs78233829 | G | G |
| rs552338116 | 18A | 18A |
| rs7380272 | T | T |
| rs11554776 | T | T |
| rs7447927 | C | C |
| rs13166214 | A | A |
| rs34703538 | A | A |
| rs7444313 | G | G |
| rs7448031 | C | C |
| rs6874798 | A | A |
| Mutation Count | 0 | 5 |

**Fig S2. The *HAQ* carriers had more generations than the *H232* carriers during the out-of-Africa migration. A-B.** A network of *HAQ* (all 22 haplotypes) and *H232* (all 12 haplotypes) found in 1kgp+HGDP individuals was constructed using Popart (popart.otago.ac.nz) by the TCS network method. The number on the line indicates the number of mutations separating the adjacent haplotypes. **C**. The population frequency and genetic diversity of the 18.5 kb *STING* haplotype were analyzed in *HAQ* individuals from the Mexican and Peruvian groups in the 1kgp population. **D.** Comparison of the 18.5 kb *STING* haplotype consisting of 19 SNPs in the ancestral *HAQ-1* haplotype and the distant, NAM-specific *HAQ-18* haplotype.

A

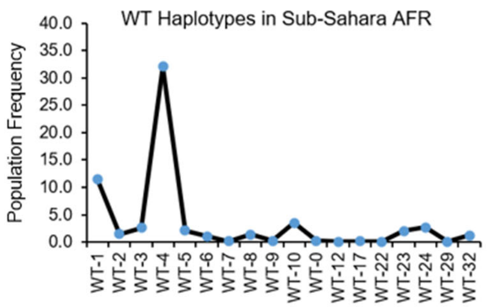


B

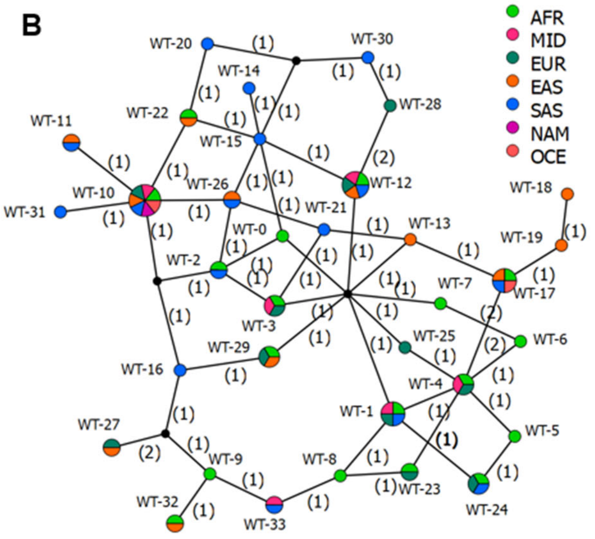


C

18.5kb; *SMIM33*; *TMEM173*

| WT Haplotypes | WT-17 (AFR) | WT-18 (EAS) | WT Haplotypes | WT-12 (AFR) | WT-30 (SAS) |
|---|---|---|---|---|---|
| rs7380062 | C | C | rs7380062 | C | T |
| rs73257323 | G | A | rs73257323 | G | A |
| rs13181561 | G | G | rs13181561 | G | G |
| rs13153461 | G | G | rs13153461 | G | G |
| rs55792153 | C | C | rs55792153 | A | C |
| rs73257329 | C | C | rs73257329 | C | C |
| rs7380824 | C | C | rs7380824 | C | C |
| rs75746446 | T | T | rs75746446 | T | T |
| rs1131769 | C | C | rs1131769 | C | C |
| rs78233829 | C | C | rs78233829 | C | C |
| rs552338116 | 21A | 18A | rs552338116 | 21A | 21A |
| rs7380272 | C | C | rs7380272 | C | C |
| rs11554776 | C | C | rs11554776 | C | C |
| rs7447927 | C | C | rs7447927 | G | G |
| rs13166214 | A | A | rs13166214 | G | G |
| rs34703538 | - | - | rs34703538 | - | - |
| rs7444313 | A | A | rs7444313 | A | A |
| rs7448031 | T | T | rs7448031 | T | T |
| rs6874798 | G | G | rs6874798 | G | G |
| Mutation Number | 0 | 2 | Mutation Number | 0 | 3 |

**Figure S3. *WT* carriers had similar numbers of generations as the *H232* carriers. A.** The population frequency and genetic diversity of the 18.5 kb *STING* haplotype were analyzed in WT individuals using 19 SNPs in 12 AFR groups in 1kgp+HGDP database. **B**. A network of WT (all 33 haplotypes) haplotypes found in 1kgp+HGDP individuals was constructed using Popart (popart.otago.ac.nz) by the TCS network method. **C**. Comparing the 18.5 kb *STING* haplotype consisting of 19 SNPs in the ancestral WT-17, WT-12 with the distant WT-18, WT-30 in non-Africans.

| HAQ Haplotypes | HAQ-1 | HAQ-2 | HAQ-3 | HAQ-4 | HAQ-5 | HAQ-6 | HAQ-7 | HAQ-8 | HAQ-9 | HAQ-10 | HAQ-11 | HAQ-12 | HAQ-13 | HAQ-14 | HAQ-15 | HAQ-16 | HAQ-17 | HAQ-18 | HAQ-19 | HAQ-20 | HAQ-21 | HAQ-22 |
|---|---|---|---|---|---|---|---|---|---|---|---|---|---|---|---|---|---|---|---|---|---|---|
| rs7380062 | T | C | C | T | T | C | T | C | C | C | T | C | T | C | T | T | C | C | T | C | T | C |
| rs73257323 | A | A | A | A | A | G | A | A | G | G | A | A | A | A | A | A | A | G | G | G | A | A |
| rs13181561 | G | G | G | G | G | G | G | G | A | G | G | G | G | G | G | G | G | A | G | A | G | G |
| rs13153461 | G | G | G | G | G | G | G | G | G | G | G | G | G | G | A | G | G | A | G | G | G | G |
| rs55792153 | A | A | C | C | A | A | A | C | C | C | A | A | A | C | A | A | C | C | A | A | A | C |
| rs73257329 | G | G | G | G | G | G | G | G | G | G | G | G | G | G | G | G | G | G | G | G | G | G |
| rs7380824 | T | T | T | T | T | T | T | T | T | T | T | T | T | T | T | T | T | T | T | T | T | T |
| rs75746446 | C | T | T | C | C | C | T | C | T | T | C | C | C | T | C | C | T | C | C | C | C | T |
| rs1131769 | C | C | C | C | C | C | C | C | C | C | C | C | C | C | C | C | C | C | C | C | C | C |
| rs78233829 | G | G | G | G | G | G | G | G | G | G | G | G | G | G | G | G | G | G | G | G | G | G |
| rs552338116 | 18A | 18A | 18A | 18A | 18A | 18A | 18A | 18A | 18A | 18A | 18A | 18A | 18A | 18A | 18A | 18A | 18A | 18A | 18A | 18A | 18A | 18A |
| rs7380272 | T | T | T | T | T | T | T | T | T | T | T | T | T | T | T | T | T | T | T | T | T | T |
| rs11554776 | T | T | T | T | T | T | T | T | T | T | T | T | T | T | T | T | T | T | T | T | T | T |
| rs7447927 | C | C | C | C | C | C | C | C | C | C | C | C | C | C | C | C | C | C | C | C | C | C |
| rs13166214 | A | A | A | A | A | A | A | A | A | A | A | A | A | A | A | A | A | A | A | A | G | A |
| rs34703538 | A | A | A | A | A | A | A | A | A | A | - | A | A | - | A | A | A | A | A | A | - | A |
| rs7444313 | G | G | G | G | G | G | G | G | G | G | G | G | G | G | G | A | G | G | G | G | A | G |
| rs7448031 | C | C | C | C | T | C | C | C | C | C | T | C | T | T | C | T | C | C | C | C | T | T |
| rs6874798 | A | A | A | A | G | A | A | A | A | A | G | A | A | G | A | G | G | A | A | A | G | G |
| Mutations Counts | 2 | 0 | 1 | 3 | 4 | 2 | 1 | 2 | 3 | 2 | 5 | 1 | 3 | 4 | 3 | 5 | 2 | 5 | 3 | 3 | 7 | 1 |

**Table S1. *HAQ* Haplotypes in Anatomically Modern Humans.** *HAQ* haplotypes were extracted from 1KGP+HGDP individuals (74 groups, ~4,000 individuals) based on a 18.5kb long genomic region consisting of 19 SNPs. The red SNPs mark the R293Q-G230A-R71H in the *HAQ*, *AQ* alleles.

| AQ Haplotypes | AQ-1 | AQ-2 | AQ-3 | AQ-4 | AQ-5 | AQ-6 | AQ-7 | AQ-8 | AQ-9/STU | AQ-10 | AQ-11/IBS | AQ-12 | AQ-13 | AQ-14/BEB | AQ-15/PLA | AQ-16 | AQ-17 | AQ-18 | AQ-19 | AQ-20/JPT | AQ-21/CHB | AQ-22/CHB2 |
|---|---|---|---|---|---|---|---|---|---|---|---|---|---|---|---|---|---|---|---|---|---|---|
| rs7380062 | C | C | C | C | C | C | C | C | C | C | C | C | C | C | C | C | C | C | C | T | T | T |
| rs73257323 | A | G | G | G | A | A | A | G | G | G | G | G | G | G | G | A | G | A | G | A | A | A |
| rs13181561 | G | A | A | A | A | A | A | A | A | A | G | A | A | A | G | A | G | G | A | G | G | G |
| rs13153461 | G | G | G | G | G | A | G | A | G | G | G | G | G | G | G | G | G | G | G | G | G | G |
| rs55792153 | A | A | A | A | A | C | C | C | A | C | A | A | C | A | A | C | A | A | A | A | A | A |
| rs73257329 | G | C | C | G | G | G | G | C | G | C | C | C | G | G | G | G | G | C | C | G | G | G |
| rs7380824 | T | T | T | T | T | T | T | T | T | T | T | T | T | T | C | T | T | T | T | T | T | T |
| rs75746446 | T | T | T | T | T | T | T | T | C | T | T | T | T | T | T | T | T | T | T | C | C | C |
| rs1131769 | C | C | C | C | C | C | C | C | C | C | C | C | C | C | C | C | C | C | C | C | C | C |
| rs78233829 | G | G | G | G | G | G | G | G | G | G | G | G | G | G | G | G | G | G | G | G | G | G |
| rs552338116 | 18A | 18A | 18A | 18A | 18A | 18A | 18A | 18A | 18A | 18A | 18A | 18A | 18A | 18A | 18A | 18A | 18A | 18A | 18A | 21A | 21A | 21A |
| rs7380272 | T | T | T | T | T | T | T | T | C | T | T | T | T | C | T | T | T | T | T | C | C | C |
| rs11554776 | C | C | C | C | C | C | C | C | C | C | C | C | C | C | C | C | C | C | C | C | C | C |
| rs7447927 | C | C | C | C | C | G | C | G | C | G | C | C | C | C | C | G | C | C | C | C | C | C |
| rs13166214 | A | A | A | A | A | G | G | G | A | G | A | A | G | A | A | G | A | A | G | A | A | A |
| rs34703538 | A | - | A | A | A | A | A | A | - | - | - | - | A | - | A | A | A | - | A | A | A | - |
| rs7444313 | G | G | G | G | G | A | G | A | G | A | G | G | G | G | G | G | G | G | G | G | G | G |
| rs7448031 | C | T | C | C | C | C | C | C | C | T | T | C | C | C | C | T | C | T | C | C | C | T |
| rs6874798 | A | G | A | A | A | A | A | A | A | G | G | G | A | A | A | G | A | G | A | A | G | G |

**Table S2.** ***AQ*** **Haplotypes in Anatomically Modern Humans**. *AQ* haplotypes were extracted from 1KGP+HGDP individuals (74 groups, ~4,000 individuals) based on a 18.5kb long genomic region consisting of 19 SNPs. The red SNPs mark the R293Q-G230A-R71H in the *HAQ*, *AQ* alleles.

.

| H232 Haplotypes | **H232-1** | H232-2 | H232-3 | H232-4 | H232-5 | H232-6 | H232-7 | H232-8 | H232-9 | H232-10 | H232-11 | H232-12 |
|---|---|---|---|---|---|---|---|---|---|---|---|---|
| rs7380062 | C | T | C | C | C | C | C | C | T | T | C | C |
| *rs73257323* | G | A | G | G | G | G | G | G | A | A | G | A |
| rs13181561 | G | G | G | A | G | G | A | G | G | G | G | G |
| rs13153461 | G | G | G | A | G | G | G | G | G | G | G | G |
| rs55792153 | A | A | A | C | A | C | A | A | C | A | A | A |
| *rs73257329* | C | C | C | C | C | C | C | C | C | C | C | C |
| *rs7380824* | C | C | C | C | C | C | C | C | C | C | C | C |
| rs75746446 | T | T | C | T | T | T | T | T | T | T | C | T |
| rs1131769 | T | T | T | T | T | T | T | T | T | T | T | T |
| *rs78233829* | C | C | C | C | C | C | C | C | C | C | C | C |
| *rs55233816* | 21A | 21A | 21A | 21A | 21A | 21A | 21A | 21A | 21A | 21A | 21A | 21A |
| *rs7380272* | C | C | C | C | C | C | C | C | C | C | C | C |
| *rs11554776* | C | C | C | C | C | C | C | C | C | C | C | C |
| rs7447927 | C | C | C | C | C | C | C | C | C | C | C | C |
| rs13166214 | A | A | A | A | A | A | A | A | A | A | A | A |
| *rs34703538* | - | - | - | - | - | - | - | A | - | - | - | - |
| rs7444313 | G | G | G | G | A | G | G | G | G | G | G | G |
| rs7448031 | T | T | T | T | T | T | T | C | T | T | C | T |
| rs6874798 | G | G | G | G | G | G | G | A | G | A | G | G |
| **Mutation Counts** | 0 | 2 | 1 | 3 | 1 | 1 | 1 | 3 | 3 | 3 | 2 | 1 |

**Table S3. *H232* Haplotypes in Anatomically Modern Humans**. *H232* haplotypes were extracted from 1KGP+HGDP individuals (74 groups, ~4,000 individuals) based on a 18.5kb long genomic region consisting of 19 SNPs. The red SNPs mark the R293Q-G230A-R71H in the *HAQ*, *AQ* alleles. *H232* is identified by rs1131769.

| WT Haplotypes | WT-0 | WT-1 | WT-2 | WT-3 | WT-4 | WT-5 | WT-6 | WT-7 | WT-8 | WT-9 | wt-10 | WT-11 | WT-12 | WT-13 | WT-14 | WT-15 | WT-16 |
|---|---|---|---|---|---|---|---|---|---|---|---|---|---|---|---|---|---|
| rs7380062 | C | C | C | C | C | C | C | C | C | C | C | C | C | C | C | C | C |
| rs73257323 | G | G | G | G | G | G | G | G | G | G | G | G | G | G | G | G | G |
| rs13181561 | G | A | A | A | G | G | G | G | A | A | A | A | G | A | G | G | A |
| rs13153461 | G | G | G | G | G | G | G | G | G | G | A | A | G | G | G | G | A |
| rs55792153 | C | A | C | C | A | A | A | C | A | A | C | C | A | C | C | C | C |
| rs73257329 | C | C | C | C | C | C | C | C | C | C | C | C | C | C | C | C | C |
| rs7380824 | C | C | C | C | C | C | C | C | C | C | C | C | C | C | C | C | C |
| rs75746446 | T | T | T | T | T | T | T | T | T | T | T | C | T | T | C | T | T |
| rs1131769 | C | C | C | C | C | C | C | C | C | C | C | C | C | C | C | C | C |
| rs78233829 | C | C | C | C | C | C | C | C | C | C | C | C | C | C | C | C | C |
| rs552338116 | 21A | 21A | 21A | 21A | 21A | 21A | 18A | 18A | 21A | 21A | 21A | 21A | 21A | 21A | 21A | 21A | 21A |
| rs7380272 | C | C | C | C | C | C | C | C | C | C | C | C | C | C | C | C | C |
| rs11554776 | C | C | C | C | C | C | C | C | C | C | C | C | C | C | C | C | C |
| rs7447927 | G | C | G | C | C | C | C | C | C | C | G | G | G | C | G | G | G |
| rs13166214 | G | A | G | G | A | A | A | G | A | A | G | G | G | A | G | G | A |
| rs34703538 | - | - | - | - | - | A | - | - | - | A | - | - | - | - | - | - | - |
| rs7444313 | G | G | G | G | G | G | G | G | G | G | A | A | A | A | A | A | G |
| rs7448031 | T | T | T | T | T | T | T | T | C | C | T | T | T | T | T | T | T |
| rs6874798 | G | G | G | G | G | G | G | G | G | A | G | G | G | G | G | G | G |

| WT Haplotypes | WT-17 | WT-18 | WT-19 | WT-20 | WT-21 | WT-22 | WT-23 | WT-24 | WT-25 | WT-26 | WT-27 | WT-28 | WT-29 | WT-30 | WT-31 | WT-32 | WT-33 |
|---|---|---|---|---|---|---|---|---|---|---|---|---|---|---|---|---|---|
| rs7380062 | C | C | C | C | C | C | C | C | C | C | C | T | C | T | C | C | C |
| rs73257323 | G | A | A | A | G | G | G | G | G | G | G | A | G | A | G | G | G |
| rs13181561 | G | G | G | G | A | G | G | A | G | A | A | G | A | G | A | G | A |
| rs13153461 | G | G | G | A | G | A | G | G | G | G | A | G | A | G | A | G | G |
| rs55792153 | C | C | C | C | C | C | A | A | A | C | C | A | C | C | C | A | A |
| rs73257329 | C | C | C | C | C | C | C | C | C | C | C | C | C | C | C | C | C |
| rs7380824 | C | C | C | C | C | C | C | C | C | C | C | C | C | C | C | C | C |
| rs75746446 | T | T | T | T | T | T | T | T | T | T | T | T | T | T | T | T | T |
| rs1131769 | C | C | C | C | C | C | C | C | C | C | C | C | C | C | C | C | C |
| rs78233829 | C | C | C | C | C | C | C | C | C | C | C | C | C | C | C | C | C |
| rs552338116 | 21A | 18A | 21A | 21A | 21A | 21A | 21A | 21A | 21A | 21A | 21A | 21A | 21A | 21A | 21A | 21A | 21A |
| rs7380272 | C | C | C | C | C | C | C | C | C | C | C | C | C | C | C | C | C |
| rs11554776 | C | C | C | C | C | C | C | C | C | C | C | C | C | C | C | C | C |
| rs7447927 | C | C | C | G | C | G | C | C | C | G | G | G | C | G | G | C | C |
| rs13166214 | A | A | A | G | G | G | A | A | G | G | A | G | A | G | G | A | A |
| rs34703538 | - | - | - | - | - | - | - | A | - | - | A | - | - | - | - | A | - |
| rs7444313 | A | A | A | A | A | A | G | G | G | A | G | A | G | A | A | G | G |
| rs7448031 | T | T | T | T | T | T | C | T | T | T | C | T | T | T | T | C | C |
| rs6874798 | G | G | G | G | G | G | G | G | G | G | A | G | G | G | A | A | A |

**Table S4.** ***WT* Haplotypes in Anatomically Modern Humans**. We extracted *WT* haplotypes from 1KGP+HGDP individuals (74 groups, ~4,000 individuals) based on an 18.5kb genomic region consisting of 19 SNPs. The red SNPs mark the R293Q-G230A-R71H in the *HAQ*, AQ alleles. *H232* is identified by rs1131769.
